# Alternatively Spliced Dual-Coding Regions Contribute to the Human Gene Regulatory Program

**DOI:** 10.64898/2026.02.24.707805

**Authors:** Clément Goubert, Alexander J. Nord, Kaitlin Sawyer, Ken Youens-Clark, George Lesica, Travis J. Wheeler

## Abstract

In eukaryotes, alternative splicing allows a single gene to encode multiple protein isoforms, by conditionally using only a subset of the gene’s exons. In some cases, distinct isoforms utilize the same exon(s) in a different reading frame, thus encoding a distinct sequence of amino acids. Here, we provide a genome-wide view of such dual coding regions (DCRs) in humans. By mapping all reviewed human UniProtKB/Swiss-Prot isoforms to the human genome and tracking reading frames used by each isoform, we identified 1296 DCR-containing genes. Though it is possible for an exon to contain multiple reading frames that lead to DCRs simply due to noisy splicing, (i) mouse orthologs to human DCR genes appear to share a dual-coding nature with much greater frequency than is expected by chance and (ii) many human and mouse DCR isoforms show differential tissue-specific expression levels, suggesting a conserved functional role. DCRs are typically short (average: 95nt), confined to a single exon, and mostly appear to introduce early stop codons that lead to loss of C-terminal coding regions. At least one third of DCRs are likely to cause nonsense-mediated decay. DCR genes are not restricted to any particular functional category, suggesting that dual coding is broadly permissive rather than confined to specialized pathways. Structure prediction indicates that most amino acids produced by canonical-frame regions are involved in some secondary structural element, while non-canonical reading frames generally produce disordered peptides, supporting a model in which dual coding primarily rewires terminal regions and isoform stability rather than creating new folded domains. Our work characterizes DCRs as a fairly common byproduct of alternative splicing, sporadically co-opted and conserved in eukaryotes through evolution, contributing to gene regulation and functional diversity. We also provide web interfaces to enable visual exploration of DCR architecture and usage patterns.

**Significance statement:** More than a thousand human genes are known to harbor coding regions that are conditionally translated in alternative reading frames through alternative splicing or transcription start site selection; however, the functional relevance of these dual-coding regions remains poorly understood. We demonstrate that this dual-coding nature is conserved: most genes containing dual-coding regions in humans also show dual-coding potential in mice. We further show that the relative abundance of proteins encoded by the alternative reading frames can vary across tissues in humans and mice and that in most cases the non-canonical reading frame produces disordered peptide domains and protein truncation. We conclude that dual coding in eukaryotes is an emergent property of noisy transcription that has been repeatedly co-opted to fine-tune gene regulatory programs.

## Introduction

Alternative splicing (AS) is a common eukaryotic process that conditionally selects different combinations of exons from a gene’s exon repertoire, thereby enabling the production of diverse protein isoforms from a single gene (Marasco & Kornblihtt 2023). Alternative splicing, combined with alternative transcription start site usage (Alfonso-Gonzalez & Hilgers 2024), plays a key role in diversifying eukaryotic proteomes (de la Fuente et al. 2025).

In a less widely recognized phenomenon, the same exonic sequence can encode different amino acids when translated in alternative frames, as a consequence of splicing variation or transcription start site selection; we use the term *dual-coding region* (DCR) to label a run of one or more consecutive exons that are observed to encode isoforms using these overlapping open reading frames (ORFs). As an example, the human *INK4A/ARF locus* codes for two proteins with distinct sequences despite sharing two exons (Sharpless & DePinho 1999). The two isoforms utilize the same E2 exon (coding) and E3 exon (primarily 3’ UTR), but differ in that INK4A/α begins from a private E1α exon, while ARF/β is initiated in an alternative upstream exon E1β; the result of splicing these alternative first exons is that INK4A and ARF process E2 and E3 in different reading frames. Though both proteins play a role in tumor suppression, they have been described as components of distinct pathways (Sharpless & DePinho 1999). Another established case of a dual-coding region is GABBR1, a component of a heterodimeric G-protein coupled receptor for GABA. A C-terminal deletion in the non-canonical isoform removes the transmembrane domain (Schwarz et al. 2000), and the truncated protein may regulate functional GABBR1/GABBR2 heterodimer formation by sequestering GABBR2 away from the canonical isoform.

While early anecdotal observations were initially thought to be an oddity (Hameed et al. 2003; Ahmed et al. 2008; Sharpless & DePinho 1999; Szklarczyk et al. 2007; Kozak 2001; Klemke et al. 2001; Scorilas et al. 2001; Zhao et al. 2004; Yoshida et al. 2001; Poulin et al. 2003), the increasing throughput of molecular readouts have provided repeated evidence of the dual-coding phenomenon, with studies identifying from dozens to thousands of DCRs from genomic (Lin et al. 2011), transcriptomic (Ribrioux et al. 2008; Xu et al. 2010; Kovacs et al. 2010; Liang & Landweber 2006; Chung et al. 2007; Vanderperre et al. 2013) and proteomic evidence (Vanderperre et al. 2013; Michel et al. 2012). The total number of human genes predicted to harbor DCRs varies depending on the evidence type and the stringency of the filters used by different research teams. Proteomic studies, which may represent the most direct evidence, produce estimates of more than 1300 genes containing DCRs (Michel et al. 2012; Vanderperre et al. 2013).

Notably, DCRs also appear to be phylogenetically conserved: (Liang & Landweber 2006)) examined 97 human genes with DCRs of length >100 nucleotides, and mapped 90 of these exons to orthologous regions in chimpanzee, of which 84 are predicted to be dual coding. (Chung et al. 2007) demonstrated, using >14,000 human/mouse transcript alignments, that DCRs larger than 500bp are extremely unlikely by chance, yet they identified 149 secondary ORFs conserved between human and mouse, suggesting underlying functionality. (Vanderperre et al. 2013) later constructed a catalog of thousands of predicted DCRs based on mature mRNA across eukaryotes, detected the peptide product of more than 1000 DCRs by mass spectrometry, and confirmed that homologs of thousands of predicted DCRs in human genes are observed in other vertebrates (chimpanzee, mouse, and cow), with hundreds observed in invertebrates (*Caenorhabditis elegans*, *Drosophila melanogaster*).

Despite several studies into the existence and role of DCRs, their prevalence seems to remain underappreciated. We first encountered DCRs serendipitously, while developing a splice-aware multiple protein isoform alignment tool, MIRAGE (Nord et al. 2018). Using MIRAGE to map isoforms from UniprotKB/Swiss-Prot (UniProt Consortium 2025) to the human genome (Nord et al. 2018), we identified 1296 genes containing dual coding regions. Importantly, DCRs analyzed in this study are limited to cases associated with curated isoforms from Swiss-Prot, meaning that this is a conservative estimate of the total number of active DCRs.

With the Swiss-Prot derived set of DCRs, we quantified frame-specific expression in 53 tissues from the GTEx consortium (GTEx Consortium 2020), and found that many of them display significant tissue-specific usage profiles across GTEx tissues with no specific gene ontology enrichment, suggesting that DCRs broadly contribute to the genome regulatory program.

Following exhaustive categorization of the splicing architecture of DCRs, we found that a large majority lead to the premature termination of the gene product; in some cases, the result is that the peptide encoded in the non-canonical frame loses some structural component of the canonical isoform (removing the encoded amino acids of C-terminal exons), while many other cases lead to transcripts that are likely subject to nonsense-mediated decay (NMD). We also predicted the 3D protein structure with AlphaFold3 (Abramson et al. 2024) and demonstrated that the large majority of non-canonical DCR peptides that are not prematurely terminated are unlikely to produce stable structures.

In summary, it appears that DCRs are a byproduct of the flexibility of the genetic code, may provide an alternative means of genomic regulation or protein reconfiguration, are utilized in tissue specific frequencies, and their dual-coding nature is conserved.

## Results

### DCR Detection

We define a dual coding region (DCR) as a coding genomic region where at least two distinct peptide sequences from alternative isoforms of the same gene can be mapped, using two or three open reading frames (**Figure 1**). Typically, a DCR overlaps a single known exon, but DCRs can also extend to multiple connected exons (**Figure 1 iii**). For clarity, we will refer to as “DCR isoform” any isoform retained in our analysis that aligns to a DCR region. Contrary to previously published work related to DCRs, we refrain here from systematically attributing an “original” or “ancestral” frame: the distinction, though often relevant, requires additional comparative analyses out of the scope of this paper. However, and whenever needed for clarity, we will refer to “canonical frame” as the frame used by the isoform labelled as “-1” by Uniprot (in case the DCR is absent of the “-1” isoform, the frame status is described as “undetermined” and frames are distinguished by their number instead).

**Figure 1.**
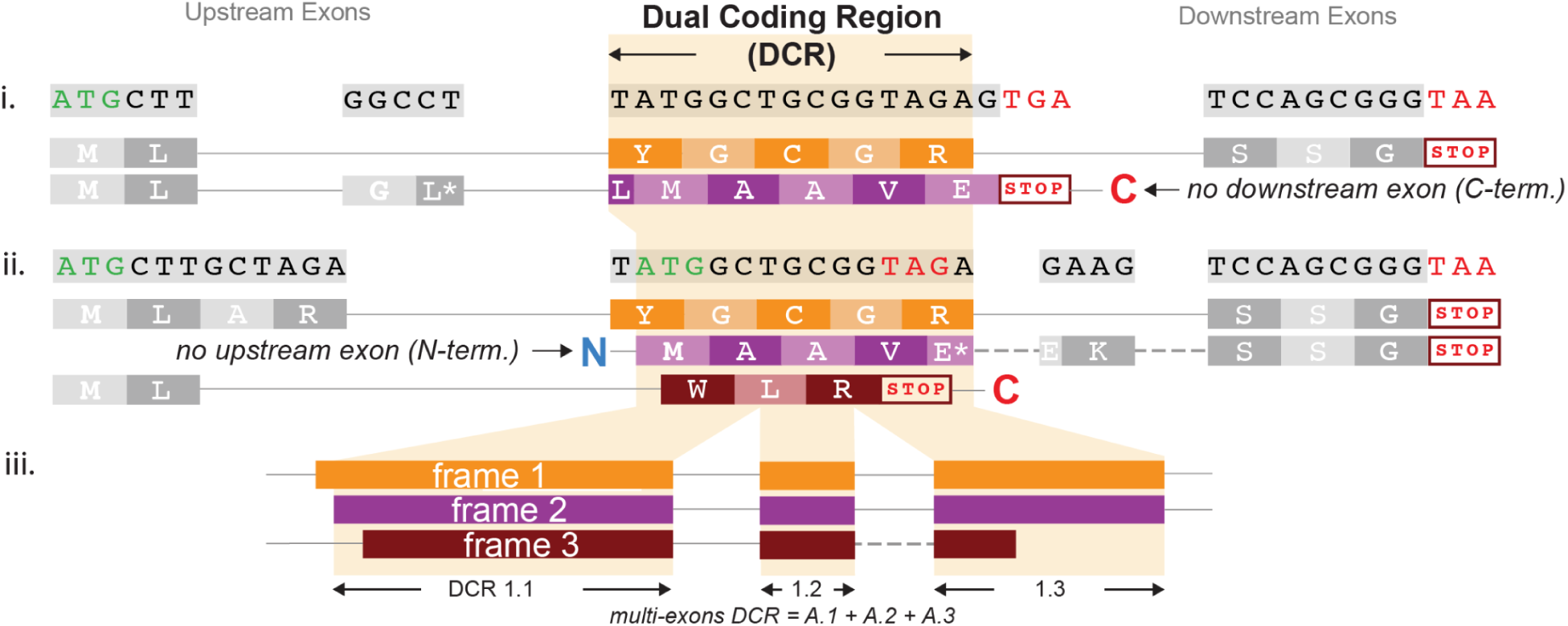
Identification of dual coding regions (DCR) from protein-to-genome mapping. A DCR represents a coding genomic region where at least two distinct peptide sequences originating from isoforms of the same gene can be mapped, using two or three open reading frames. DCRs are mainly the product of alternative splicing, and can lead to early termination of the protein (i) but can also be caused by alternative CDS initiation start (ii). While most DCRs cover a single exon, cases can include multiple successive exons (iii)

Using MIRAGE2 (Nord et al. 2018), we aligned a total of 39,714 protein isoforms from UniprotKB/Swiss-Prot to the hg38 human reference genome (GRCH38.p14). MIRAGE identified 1573 candidate genes with DCRs that were not the product of intron-retention events. Following quality filtering (see Method), 1308 DCR-containing genes were retained (Table S1). After manual inspection of edge-cases, we withdrew 24 DCRs (12 genes) from the subsequent automated bioinformatic analyses, where complex combinations of splice sites induce distinct groups of isoforms to use separate open reading frames in successive DCRs along the same gene. These special cases are discussed in a separate section of this manuscript. Unless otherwise stated, the analyses are thus performed on a set of 1407 DCRs, representing 1296 human genes (Table S1).

### Conservation and evolutionary constraints

#### Conservation of the Dual Coding status between human and mice

One way to validate that a genomic feature is functional is to observe that it is conserved. We tested this by checking if human DCRs are also dual coding in mice. Because DCRs can span across multiple exons, the 1296 DCR genes observed in humans and retained for automated analyses, can be broken down into 1407 DCRs, represented by 1741 unique genomic intervals, where an interval is a portion of an exon used by both isoforms. Of these, 1603 exons (representing 1295 human DCRs) could be mapped to the mouse mm39 assembly. Ideally, we would be able to test if these homologous mouse intervals were also observed to encode two or more isoforms in Swiss-Prot; this is not effective because isoform data for mouse is much less complete than for human (at the time of this writing, Swiss-Prot contains 52,582 human isoforms, compared with 38,032 mouse isoforms). Instead, we tested if the homologous mouse regions have DCR *potential*, meaning that they encode two reading frames over their entire length (allowing for minor splice site migration by trimming the mapped intervals by 1 codon to on both 5’ and 3’ ends). We restricted analysis to only intervals of length between 21 and 500 nucleotides. Of the 1294 human DCRs with this length constraint, 981 (80%) are also dual coding in mice **Figure 2A**. Only 3 human genes have an exon overlapping a DCR > 500 nt; of these, *GNAS* is the only one whose dual-coding potential appears to be conserved in mice (data not shown). To assess if this level of conservation is expected by chance, we compared the proportion of dual coding mice intervals with two controls. In the first control, we sampled 100,000 random intervals (≥21nt and ≤500nt) from all non-DCR mouse exons (trimming instances to match the length distribution of the DCR intervals). In the second control, we randomly shuffled codons for all mouse CDS ≥21nt and ≤500nt (N = 493,640). **Figure 2B** shows that mouse intervals orthologous to human DCRs are potentially dual coding with higher frequency than the controls in all but 6 size bins for which only a handful of DCR intervals are available.

**Figure 2.**
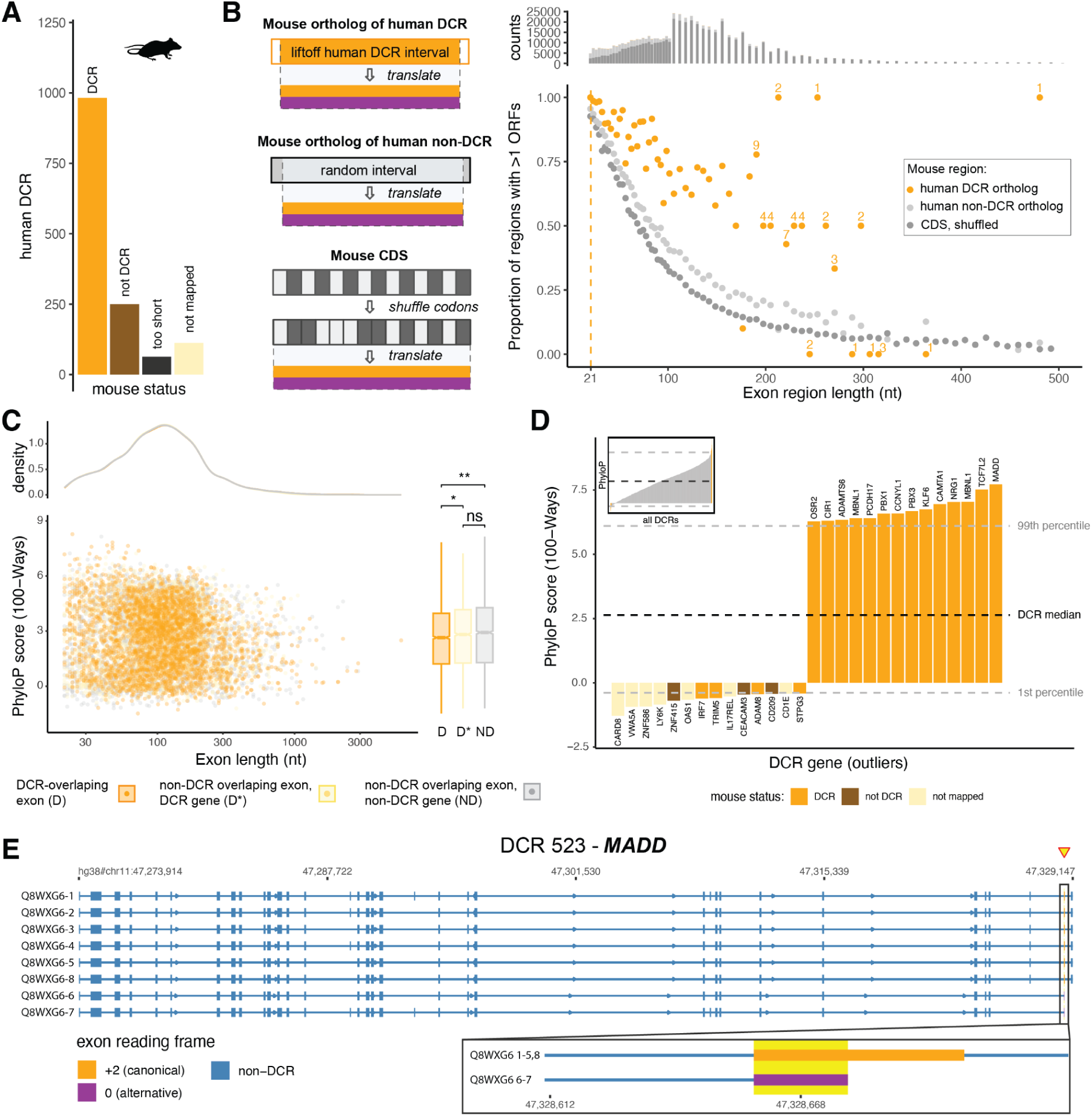
Dual Coding Regions conservation. **A.** Conservation of the dual coding status of human DCRs after LiftOver to the mouse genome mm39. The mouse region must produce at least 2 ORFs that cover the full length of the mapped sequence, allowing for at most one missing codon on either end. In cases of multi-exon DCRs, some mouse genes have dual coding potential on only some of the mapped exons; we categorized them as “not DCR”. **B.** Proportion of regions of the mouse genome (mm39) orthologous to human DCR exons (orange) or controls (greyscale), producing 2 or 3 ORFs, binned by region length and limited to regions ≤ 500bp. The orange vertical dotted line indicates the 21nt cutoff used for observed DCR intervals **C.** Distribution of the 100-ways PhyloP conservation score for human exons according to their overlap to DCR and non-DCR exons in DCR genes. **D.** Outlier human DCRs (first and last percentiles of the 100-ways PhyloP conservation score) and associated genes **E.** Gene model for *MADD*, which is the most conserved DCR gene reported in our study. Two isoforms (Q8WXG6-6 and -7) use an alternative ORF in their last exon (one-before-last in the gene model) causing dual coding and a premature end of the protein relative to the other isoforms.

#### DCR sequence constraint

We further asked whether the nucleotide sequence of DCR presented a different selection signature compared to ordinary exons. As a proxy of selection strength, we leveraged per-base phyloP (Pollard et al. 2010) scores computed for 100 vertebrates on hg38 (**Figure 2C**) available via the table browser at UCSC Genome Browser (Perez et al. 2025). A positive phyloP score indicates conservation, while negative scores suggest diversification (Pollard et al. 2010). We computed per-exon average phyloP for DCR-overlapping exons and two controls: non-DCR-overlapping exons within DCR genes, and exons in non-DCR genes. Because DCR-overlapping exons are generally smaller than the control exons, we resampled control exons to match the size distribution observed in DCRs (Figure 2C, top panel). DCR-overlapping exons are slightly (yet significantly) less conserved than both controls (Student’s t-test with Boneferroni correction: DCR vs non-DCR in DCR genes: *P* = 0.02; DCR vs non-DCR in non-DCR genes: *P* = 0.001), while there is no statistical difference between the two controls (*P* = 0.314). Taken together, the results suggest that evolutionary constraints on DCRs are globally similar to other exons, though selection may have been slightly relaxed in a subset of DCR genes.

### Characterization of human Dual Coding Regions

#### DCR architecture

We categorized the diverse DCR architectures based on the type of junctions emitted upstream and downstream of the exon(s) located in the DCR for the isoforms involved. Individual DCR architectures can be visualized online through our dedicated browser: https://cgoubert.shinyapps.io/DCR_Architecture/. The first criteria is whether the DCR is N- or C-terminal for one or more isoforms. Next, if an isoform has junctions with upstream or downstream exons we inspect two more criteria: (1) whether the boundaries of the exons overlapping the DCR region are identical (“DCR exon boundaries: same”) or different (“DCR exon boundaries: distinct”) between isoforms using different frames, and (2) whether the exon junctions connected to the DCR are identical (“linked exon boundaries: same”) or different (“linked exon boundaries: distinct”) between isoforms using different frames (**Figure 3A**). The large majority of cases (1150/1407, 81.7%) induce the C-terminal truncation of the protein for one or more isoforms. In contrast, 17.1% (241/1406) of DCRs are N-terminal for at least one isoform and only forty cases (2.8% of the total) include combinations of isoforms with either N-or C- terminal truncations at the DCR. Finally, 4% (56) of the DCRs are entirely internal, meaning that a splicing event causes an exon to be used in a non-canonical frame, then another splicing event reverts the isoform to the standard frame. Among 45 unique architectures, three involving C-truncation represent nearly two thirds (864/1407, 61.4%) of the cases. The most abundant architecture, seen in 405 DCRs, occurs when an exon is used by two isoforms that use distinct upstream exons, each entering the shared exon in a different reading frame, and resulting in one isoform that encounters a stop codon within the dual coding region (i.e. is C-terminal (**Figure 3A**, case a)). The second most abundant architecture (N = 347) shares the upstream pattern with case a, but differs in that the DCR is C-terminal for all isoforms (**Figure 3A**, case b). We noticed that genes that harbored two or more DCRs often do so because of the inclusion of exons exclusively found in groups of isoforms that share the same reading frames across DCRs. Thus, in these cases, even if an upstream DCR does not directly cause a C-terminal truncation, downstream ones often do, so that 94.8% (92/97) of these genes contain truncated isoforms due to dual coding, contributing to 82.8% (1073/1296) of all DCR genes.

**Figure 3.**
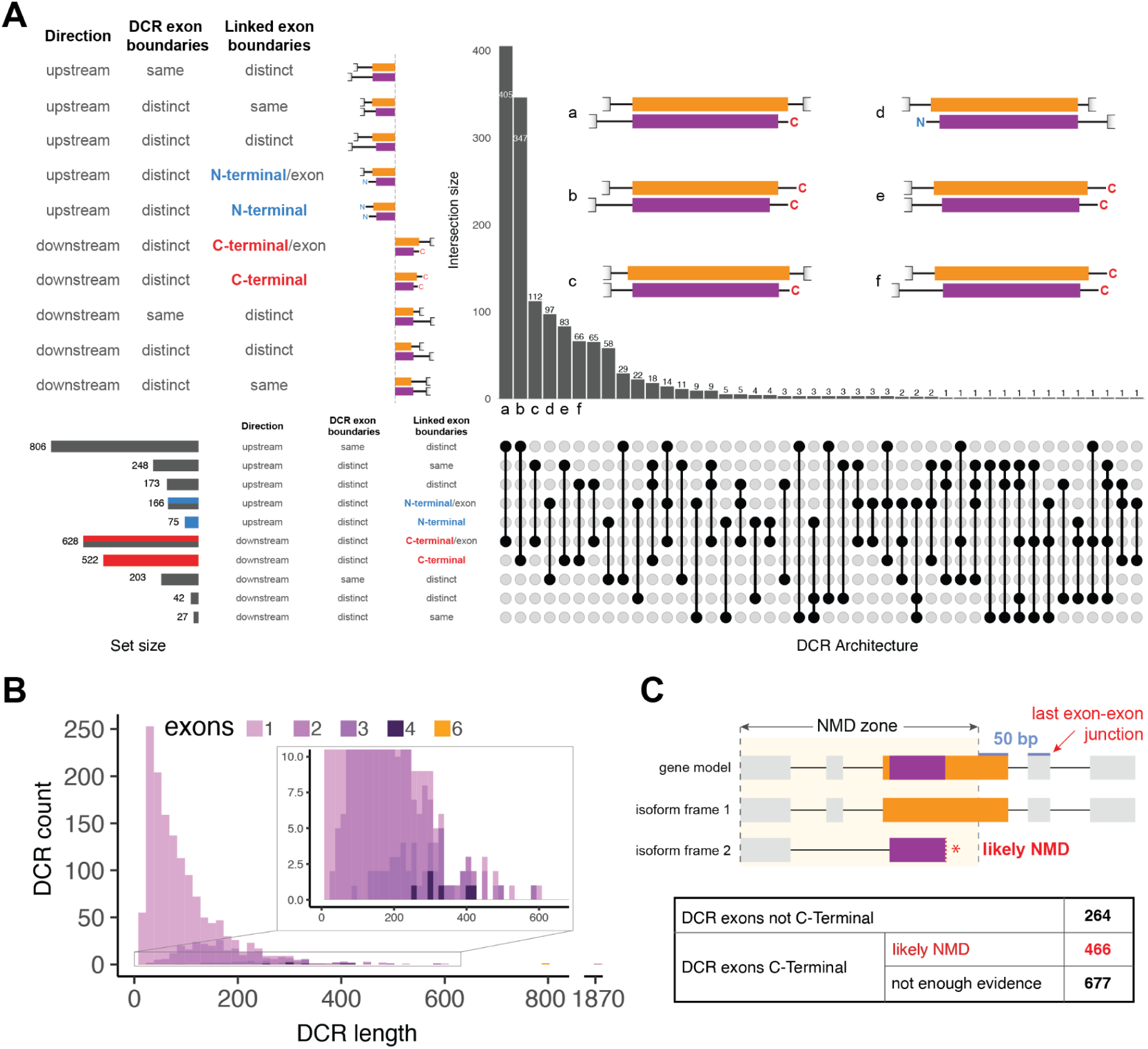
Characterization of Dual Cording Regions in the human genome. **A.** Upset plot representing the different architectures of DCRs based on their upstream and downstream exon junctions. **B.** Distribution of the DCR region lengths (nt) colored according to the total number of exons within the region. **C.** Nonsense-mediated decay susceptibility of truncated isoforms caused by DCRs. “Not enough evidence” indicates cases where the DCR lies in the penultimate exon, but NMD status cannot be determined because the 3′UTR location of isoforms is unknown (mapping is done at the protein level).

Almost all (N=1391, 99%) DCRs involve contests between two frames, while 16 (1%) involve 3 overlapping frames. The DCRs range from 20 to 1878 nucleotides in length, with a mean of 95nt±92 s.d. (**Figure 3B**). While most DCRs are confined to a single exon (N = 1131, 80%), the 276 remaining cases extend to two (N = 227, 16%) or more exons (3 exons: N = 42; 4 exons: N = 6), up to 6 exons (N = 1).

A systematic analysis of the exon-exon junctions motifs upstream and downstream DCRs did not reveal a bias from the canonical 5’-GT…AG-3’ signal (Burset et al. 2000) when DCRs are compared to regular exon-exon junctions (Figure S1).

#### Susceptibility to Nonsense-Mediated Decay

Nonsense mediated decay (NMD) is a regulatory cellular process in which a transcript containing a premature stop codon is quickly recognized and degraded. It is commonly accepted that if a stop codon occurs at least 50-55 nucleotides upstream of the final exon-exon junction, the transcript is subject to NMD (Nagy & Maquat 1998; Hug et al. 2016). DCRs encompass cases where some isoforms, using a first frame, connect to a downstream exon while others, using a second or third frame, incorporate an apparent premature stop codon (**Figure 3A,C**). Therefore, we expect that many DCR isoforms will contain a stop codon located more than 50 nt upstream of putative exon-exon junction in the expected pre-mRNA molecule. Our investigation of NMD in DCRs is limited by the fact that by directly mapping protein isoforms, we do not have access to the 3’ untranslated region of the underlying mRNA and thus do not know with certainty the location of the last exon-exon junction. Nevertheless, we can identify isoforms that are likely subject to NMD by displaying a stop codon at least 50 nt upstream of the last observed exon-exon junction. Accordingly, we estimate that 33.1% (N = 466/1407) of DCR cases may lead to isoforms subject to NMD, while 18.8% (N = 264) are predicted not to be the target of NMD due to the presence of a downstream exon, and the remainder face an undetermined fate due to unclear 3’ UTR status. Though the UniprotKB/Swiss-Prot dataset is restricted to proteins with a “reviewed” status, an analysis of the UniProt annotation codes shows that non-canonical isoforms, and in particular those flagged as potential NMD target in this study, have generally weaker evidence as reported by the ECO ontology codes (Figure S2).

### Examples of human DCR genes with high conservation score

Among the most conserved human DCR genes, we identified *MADD* (MAP kinase-activating death domain protein). Two MADD isoforms (Uniprot ID Q8WXG6-6 and Q8WXG6-7) skip over exon 30 and connect to exon 31 in a second non-canonical frame before reaching a stop codon after translating 7 amino-acids, prematurely ending the protein (**Figure 2E**). A comparison of isoforms 1 and 6 with Interproscan (Blum et al. 2025) shows that the short form trims, but does not remove, a C-terminal Pleckstrin homology (PH) domain responsible for binding signal proteins to cell lipid membranes (Cozier et al. 2004). Another conserved DCR example is *TCF7L2*, a critical transcription factor of the *Wnt* signaling pathway involved in glucose homeostasis. Isoform 2 and 4 (Uniprot ID Q9NQB0-2,3) introduce a penultimate exon 13 leading to a premature stop codon in their last exon (14), relative to isoforms 3, 7 and 13 representatives of the long (E) isoforms of this gene, that include a C-clamp motif required for DNA binding ((Weise et al. 2010), Figure S3).

#### DCRs do not appear to be associated with functional categories

Gene Ontology (GO) term enrichment analysis, performed with the PANTHER classification system (Mi et al. 2019) for the 1338 DCR genes, did not yield significant results, indicating that the phenomenon is broad and affects genes independent of their functional annotation.

#### Complex DCR architectures

Our bioinformatic classification of DCR architectures identified edge cases for 12 genes that each harbor 2 distinct but interdependent DCRs. In contrast with other genes harboring multiple DCRs, complex scenarios implicating interwoven splice site usage caused adjacent DCRs to cluster isoforms in distinct groups according to the frame used to translate the exon. The 12 genes are *CYB561D1, ENOSF1, FUT8, GIMAP6, IL23R, LARP1B, MAEA, METTL26, MFSD1, NRG2, RNF170, UBAC2*;their architectures are included in the visualizer available at https://cgoubert.shinyapps.io/DCR_Architecture/

### DCR differential isoform abundance across tissues

#### A subset of DCRs display significant frame-by-tissue specificity across GTEx tissues

We tested whether the reading frame used by DCR isoforms shows tissue-specificity, as a proxy of functional relevance. We quantified isoform usage based on the abundance of exon-exon junctions (EEJ) entering and exiting DCRs across 53 GTEx tissues (Supplementary Data 1, see Methods). We retained EEJ with an average of 2 reads per sample in at least 2 tissues and computed pairwise Peasron’s correlation between tissues (Figure S4).

We implemented a simple statistical test based on the difference in frame usage ratio between tissues (tissue x frame interaction), followed by a screen to identify cases in which different tissues show significantly-different patterns of DCR isoform usage (see Methods). We identified 108 DCRs with significant tissue x frame interaction (min. Boneferroni-corrected *P*-value < 0.05 between two tissues), of which, 65 were classified as disordinal (with one tissue preferring an isoform with one reading frame, and a different tissue preferring an isoform with a different frame; Table S2, **Figure 4**). A manual review of the candidate DCRs showed a clear brain-specific usage pattern in multiple candidates, notably for the DCR genes *MARK4*, *MADD*, *CAMKK2*, *ATP2B1*, *PLCB4*, *STX1A*, *AMPD2* and *GABBR1*. An interactive visualization of these results, including per-DCR tissue x frame expression levels and a representation of the DCR with isoforms mapping in the genome coordinate system at https://cgoubert.shinyapps.io/DCR_Explorer.

**Figure 4.**
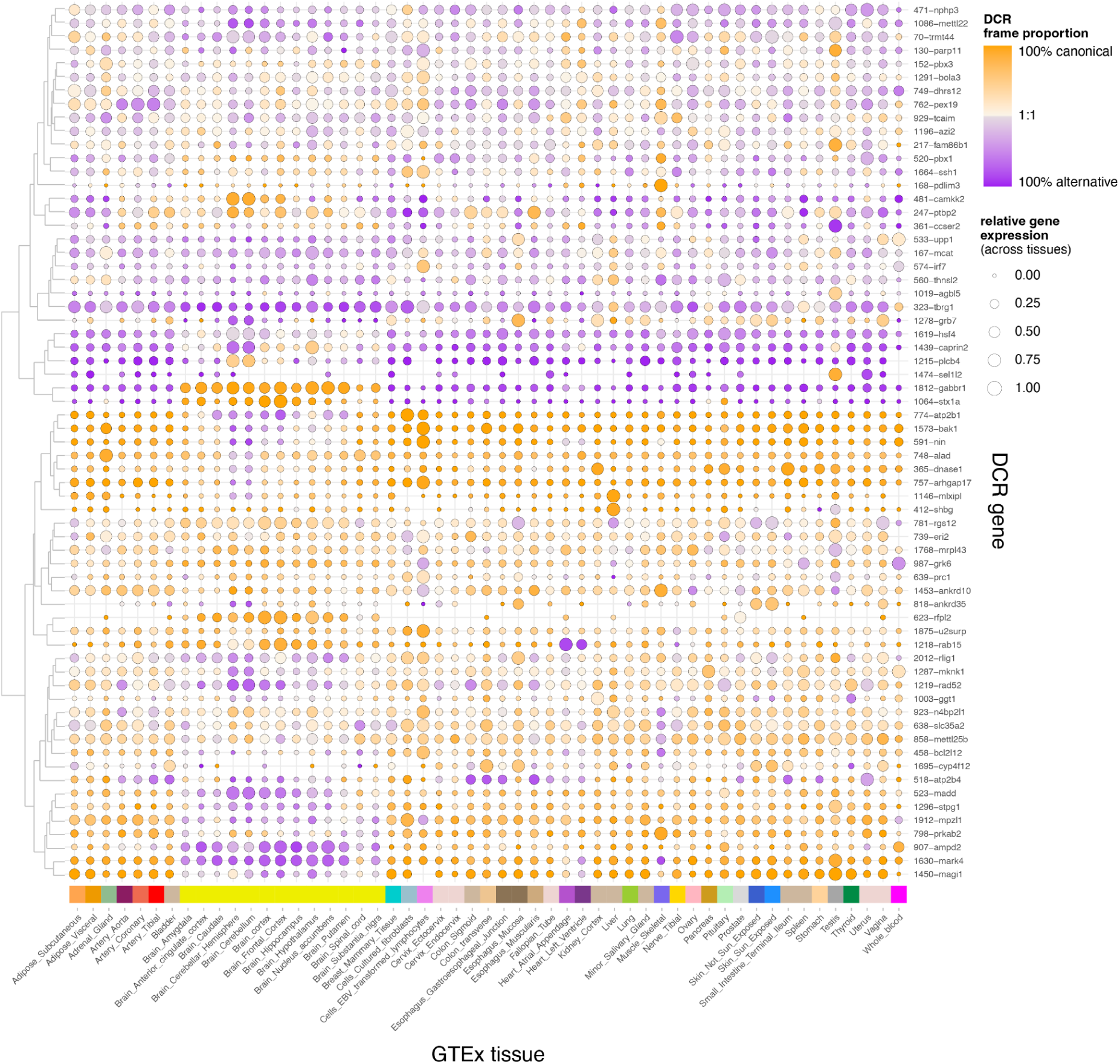
Differential frame usage at 65 candidate DCRs in 53 GTEx tissues. Each row represents a candidate DCR satisfying the thresholds of the statistical analysis, while each column represents a tissue from the GTEx dataset. Circle size is proportional to the relative gene expression, normalized across all tissues. The color shade represents the proportion of reads originating from isoforms using canonical (orange) or alternate (purple) reading frames in the DCR.

#### DCR frame usage tissue-specificity agrees with gene expression patterns

While the final quantified dataset only comprised 1263 genes, DCR’s junction expression showed correlation between similar tissues, particularly in the brain as well as tissues rich in smooth muscles (such as arteries, uterus, the gastroesophageal junction of the esophagus and colon sigmoid), stromal and fat rich tissues (adipose and mammary), epithelial-rich tissues (vagina and skin), as well as grouping together liver and pancreas (digestion and nutrient processing) (Figure S4). Though these results may simply recapitulate tissue-specific gene expression rather than DCR frame-usage specificity, it confirms the permissivity of DCRs, suggested by the lack of GO-term enrichment for the DCR gene set.

#### Mouse DCR orthologs also show evidence of tissue-specific frame usage

We reasoned that human DCR genes with evidence of frame-by-tissue interaction may also exhibit such properties in mouse orthologs. We leveraged the transcriptome profiling of multiple mouse tissues performed by (Li et al. 2017) and quantified frame-specific EEJs for 12 mouse orthologs of human DCR genes (see methods). Ten out of the 12 genes screened showed evidence of dual frame usage by alternative isoforms in mice, with 7 showing frame switching pattern between tissues. For some of these genes, such as *MADD* and *ATP2B4,* the frame switching pattern between tissues was similar to humans across the majority of tissues tested (**Figure 5**).

**Figure 5.**
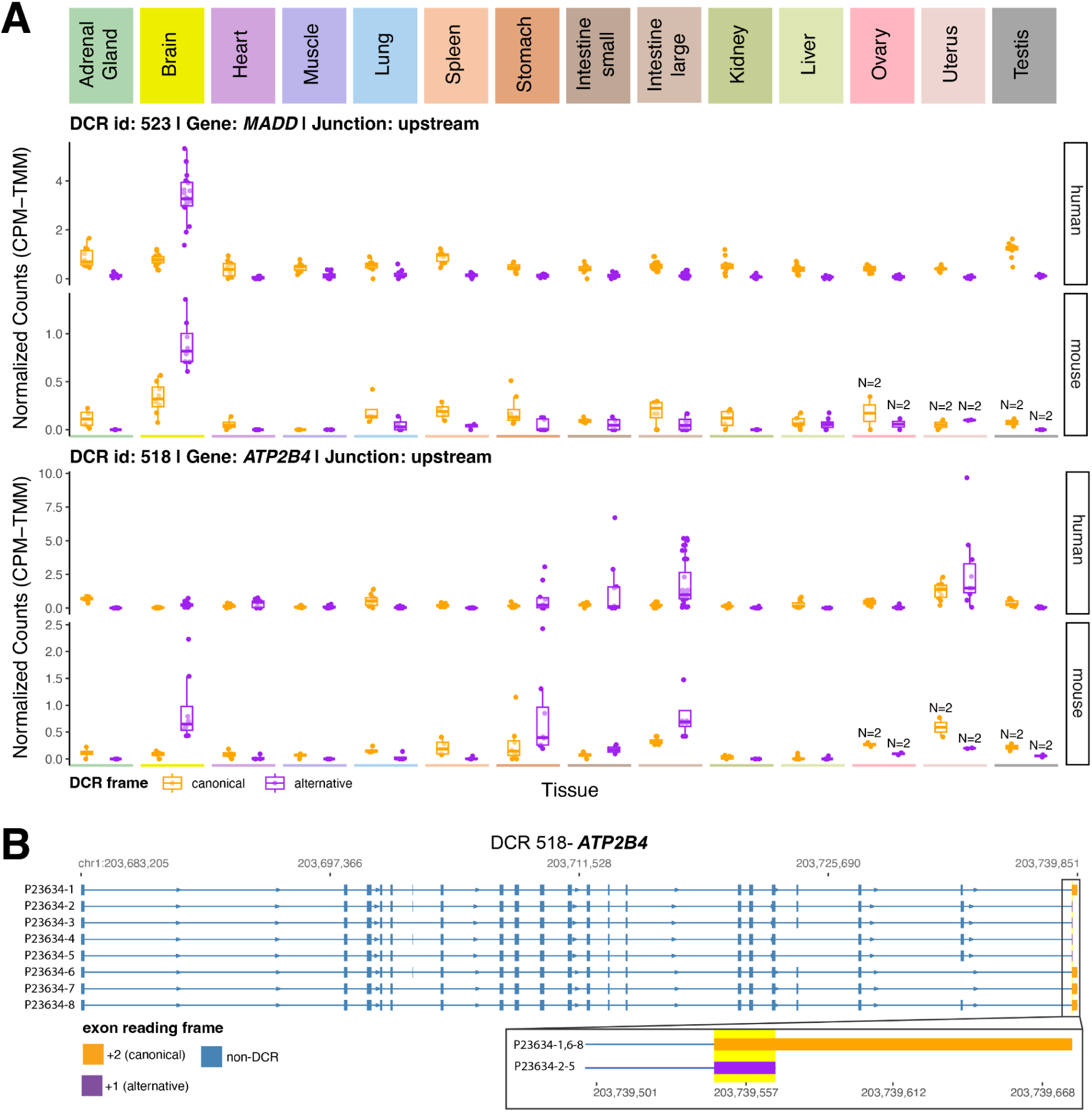
Differential frame usage at DCR in human and mouse. **A.** Normalized counts of upstream EEJ leading to DCR for two orthologous genes in human and mouse. For each gene, the counts are separated according to tissues (columns) in each species (rows). Within each tissue, counts are separated according to the frame used in the DCR: canonical (orange) or alternative (purple). Boxplots are overlaid on top of individual tissue replicates. Top: *MADD*, bottom: *ATP2B4*. **B.** Gene model from protein isoforms sequences of the human gene *ATP2B4*.

### Non-canonical DCR peptides show limited evidence of 3D structure

To gain insight into the function of the distinct peptides encoded by isoforms using a non-canonical frame at DCRs, we performed protein structure predictions for 56 pairs of isoforms representative of the canonical or alternative frame for a subset of DCR. We focused on DCRs that do not occur due to an alternate start and do not cause a premature end to the protein isoform. In other words, we looked at DCRs where each alternate frame, carried by distinct isoforms, connects to an exon upstream and downstream. This approach essentially retains cases where the DCR induces a peptide switch within the protein sequence. We used Alphafold3 (Abramson et al. 2024) to predict each isoform structure and extract per-amino acid plDDT in the DCR region. We also registered the predicted secondary structure classification of each amino acid (alpha helix, beta ladder, etc…) according to DSSP (Hekkelman et al. 2025). We found that DCR peptides of isoforms using the non-canonical frame were overwhelmingly lacking evidence of structure in comparison to canonical isoforms (**Figure 6A**), suggesting that DCRs are most likely to ablate function rather than providing a novel biochemical property. As an example, the DCR region of the gene *DAGLB* (**Figure 6B**) is predicted to encode a peptide contained in an alpha helix with strong support from the model (**Figure 6B** i. and ii., Alphafold3, plDDT from >70 to >90 range), while the alternate frame of the DCR is predicted to encode a disordered region and fragments of alpha helices but with very low support (**Figure 6B** iii. and iv., Alphafold 3, plDDT < 50).

**Figure 6.**
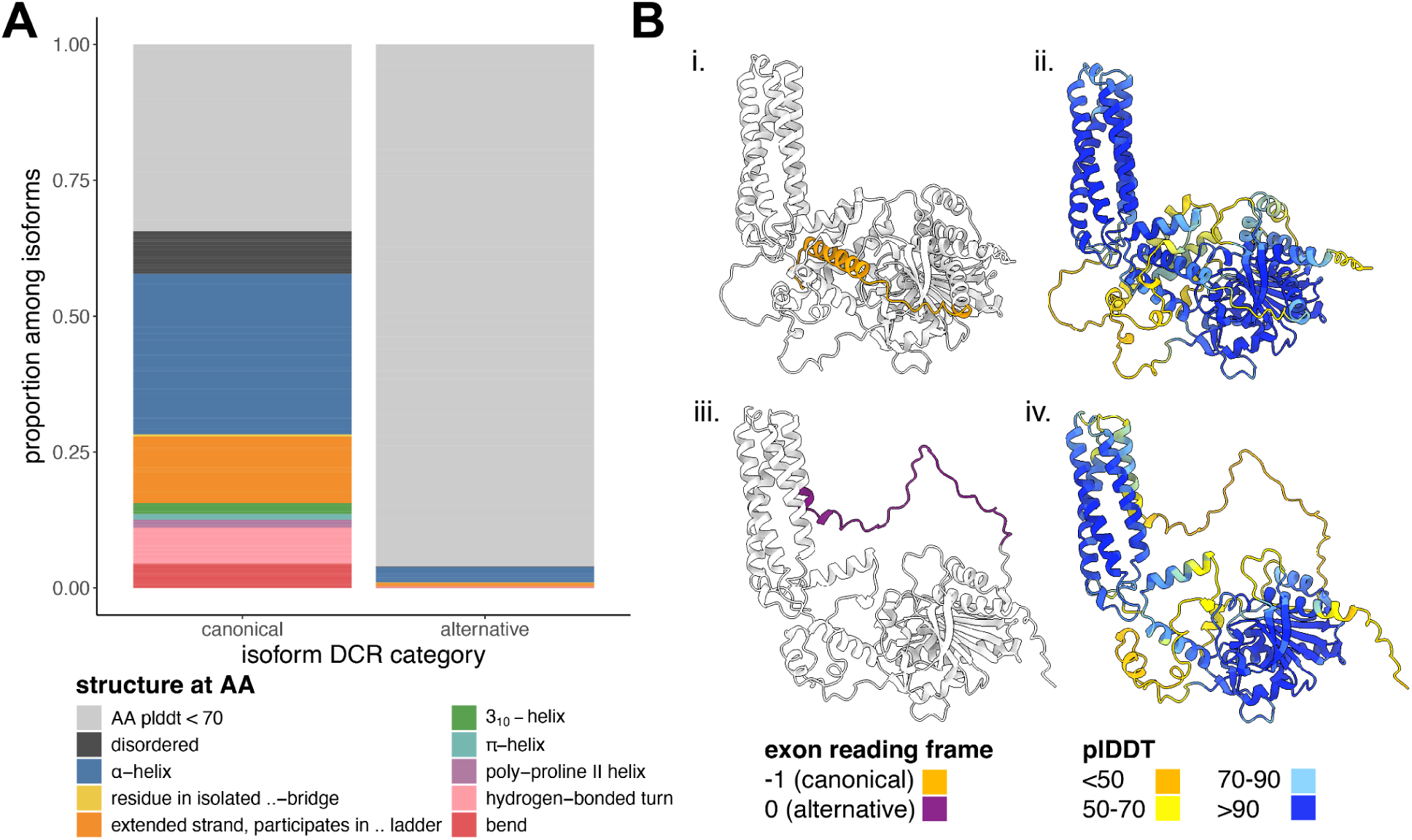
DCR structure prediction. **A.** Structure prediction at the peptide regions encoded in a subset of DCR isoforms. **B.** Alphafold3 structure prediction for two isoforms of *DAGLB* (DCR 767) representative of the canonical (i., ii.) and alternative (iii., iv.) frames in the DCR.

## Discussion

Dual coding regions (DCRs) are a puzzling feature in eukaryotic genomes, repeatedly observed in mammals, with dual coding orthologs of human DCRs found as far as in invertebrates and even yeast. Though previous efforts have characterized their prevalence and evolution, few studies have focused on their functional relevance (Kovacs et al. 2010; Vanderperre et al. 2013). Here, we have addressed this gap by leveraging 39,714 reviewed human protein isoforms of the UniprotKB/Swiss-Prot database, combined with transcriptome profiling in 53 human tissues and additional investigations for 14 tissues of the mouse transcriptomic BodyMap, and we further explored the influence of DCRs on the predicted structure of human proteins.

We identified 1296 genes containing 1407 dual coding regions of an average length of 95 bp, ranging from 20 to 1,878 bp. The extent of genes identified in the literature as containing DCRs is widely variable, due to differences in data sources (conserved CDS, transcripts or protein evidence), timeline (reports spanning nearly 30 years of research in molecular biology) and bioinformatic thresholds. Nevertheless, our count is very close to the total of 1,310 distinct genes gathered from protein evidence summed over two studies (Michel et al. 2012; Vanderperre et al. 2013). From this comparison, we identified 75 novel DCR genes, suggesting that these reports, including ours, are underestimates (Figure S5). Such results are expected: the UniprotKB/Swiss-Prot dataset only includes proteins with reviewed evidence, while data from (Vanderperre et al. 2013) was compiled from an incomplete set of tissues, and required refSeq overlap and mass spectrometry detectability.

The large majority of DCRs involve changes in the final exons of the genes. Because non-canonical final exons typically encode disordered C-terminal tails and distinct 3′UTRs, isoforms that differ only in their last exon are most often expected to diverge in subcellular localization, interaction partners and signaling output, protein stability, and post-transcriptional regulation, rather than in their core catalytic activity (Mayr 2019, 2017; Preussner et al. 2020; Rodriguez et al. 2020; Ravichandran et al. 2024). Furthermore, at least 33% of our candidates were predicted to produce isoforms susceptible to nonsense mediated decay (NMD) under the 50bp rule (Nagy & Maquat 1998). NMD is generally seen as a means to reduce transcriptional noise, so observing DCR-derived products as NMD targets may suggest that dual coding is only an accidental byproduct of alternative splicing (AS), and to a lesser extent, the selection of alternative transcription start site. Kovacs et al. (2010), downplayed this possibility, initially reporting that only 5% of DCRs lead to NMD, but only screened 62 cases. With the caveat that UniprotKB/Swiss-Prot isoforms flagged for NMD tend to have weaker type of evidence support (Figure S2), we hypothesize that dual-coding was, in these particular cases, co-opted as a way to silence gene expression in a tissue-specific manner, a mechanism sometimes referred as regulated unproductive splicing and translation (RUST) (Lewis et al. 2003). In fact, among 28 DCR genes that show frame-by-tissue differential expression and for which at least one isoform is susceptible to NMD, four have been independently shown to exhibit RUST-like behavior, including *PTBP2, BAK1, GABBR1, and ARHGAP17 (Vuong et al. 2022; Ling et al. 2016; Lee et al. 2023; Makeyev et al. 2007; Mauger & Scheiffele 2017; Linares et al. 2015; Najar et al. 2025)*.

It is worth mentioning that our transcriptomic screening represents a narrow view of the potential functionality of DCR genes. To clearly distinguish biological signal from transcriptional noise, we deliberately applied stringent filters on the relative expression of each frame, and also required disordinal interaction between frames and tissues. It remains likely that differential frame usage occurs between tissues in more moderate ways, however, we argue that testing such cases would require a stronger statistical framework using isoform-level quantification. Furthermore, the GTEx samples only represent a set of conditions across healthy adult tissues (GTEx Consortium 2020). Investigating differential frame usage of DCRs across developmental stages or health conditions, in particular in the context of cancer and aging, may further garner insight into the biology of DCRs.

In their analysis, (Kovacs et al. 2010) showed bioinformatically that in eukaryotes, maintaining two overlapping functional ORFs can only be achieved if the derived peptide is more intrinsically disordered than its canonical counterpart, owing to the fact that eukaryotic genomes lack both the spatial constraints and the large effective population sizes of pathogenic prokaryotes that would permit the evolution of two completely distinct, well-folded proteins from the same sequence. In addition, the authors suspected but could not demonstrate that AS was responsible for most of their observed DCRs, citing correlation between AS and structural disorder (Romero et al. 2006). Our comprehensive review of the UniprotKB/Swiss-Prot dataset unambiguously confirms that AS is the driving mechanism behind eukaryotic DCRs.

Furthermore, our structural analysis of the DCRs-encoded peptides with Alphafold3 concurs with the findings of Kovacs et al. (2010): focusing on DCRs affecting internal regions of the genes, we found that the amino-acids encoded by the alternative frame virtually never met the trusted threshold indicative of a 3D structure (plDDT > 70). One important caveat is that AlphaFold3 structure prediction depends on sequence alignments involving multiple orthologous sequences, and non-canonical isoforms are likely to gather fewer (or possibly no) such sequences to aid in templating; as a result, it is possible that disorder prediction in non-canonical peptides may be overestimated.

If we can conclusively assert that DCRs are primarily a fortuitous consequence of AS, sporadically selected for its functionality across eukaryotes, we can hypothesize that such atypical arrangement of coding sequence could only evolve under certain circumstances. We found that the nucleotide sequence of exons overlapping DCRs are slightly (but significantly) less conserved than regular exons. Indeed, an increased tolerance for substitution might be a requirement toward evolving a secondary open reading frame. (Liang & Landweber 2006) for instance reported that derived secondary reading frames (determined to have evolved later) had increased amino acid substitution rate between human and chimpanzee relative to the ancestral reading frames. (Vanderperre et al. 2013) later confirmed these results, expanding their observation to other mammals, amphibians, invertebrates and yeast. Nevertheless, the conservation score of DCRs vary widely and still include very conserved genes such as (i) *MADD*, which regulates apoptotic processes, (ii) *TCF7L2* (Transcription factor 7-like 2), involved in many signal transduction pathways, and (iii) *MBNL1* (Muscleblind-like 1) which, intriguingly, is itself a key regulator of alternative splicing. Together, these results suggest that while relaxed selective pressures may favor dual coding, the overall plasticity of the genetic code allows a broad collection of genes to diversify their isoform repertoire via dual coding.

## Conclusion

Our analysis of 39,714 reviewed proteins isoforms of the UniprotKB/Swiss-Prot database suggest that dual coding is principally the product of alternative splicing, sporadically co-opted and phylogenetically conserved throughout the evolution of eukaryotes, enabling diversification of the proteomic repertoire. DCRs principally affect the C-terminal regions of proteins (though N-terminal and internal variation exist), replacing functional domains with disordered or truncated peptides. In consequence, the alternative isoforms may provide functional diversity through the regulation of existing functions, for instance *via* unproductive splicing, rather than providing novel functional sites or domains.

## Limitations

Our survey is conservative by design. By restricting ourselves to manually curated UniprotKB/Swiss-Prot isoforms, healthy adult GTEx tissues and a limited mouse panel, we almost certainly miss context-specific, lowly expressed, or unreviewed overlapping ORFs. Our detection of frame-by-tissue interactions uses stringent thresholds, so more subtle shifts in frame usage are likely undercalled. Finally, our structural conclusions rely on AlphaFold3 predictions for internal DCRs, which may lack power in poorly aligned regions (which are likely to occur for under-explored splicing variants); thus, we are biased toward cases with clearer structural signals and likely underestimate the full structural and regulatory diversity of dual coding regions. We note that the definition of canonical frame, based on the UniprotKB/Swiss-Prot nomenclature, does not always reflect the frame preferentially used across most GTEx tissues (e.g. DCRs 323,1439, 1812, 1064). Perhaps a more meaningful description involves the classification of the frames as ancestral vs derived. Such a framework would require dedicated phylogenetic reconstruction which was out of the scope for this work.

## Methods

Unless otherwise stated, statistical analyses and figures were produced with R version 4.3.1.(R Core Team 2023) and Python 3.11. When necessary, relevant packages are indicated.

Datasets, custom programs and analysis scripts are publicly available at https://github.com/TravisWheelerLab/DCR-project.

### DCR discovery

Dual coding regions (DCRs) were reported by first mapping the protein sequences of 39,714 human isoforms present in the curated set UniprotKB/Swiss-Prot (UniProt Consortium 2025). We downloaded isoforms from https://ftp.uniprot.org/pub/databases/uniprot/current_release/knowledgebase/complete/uniprot_sprot.fasta.gz, last accessed on May 27, 2025 (release-2025_01), then mapped to human hg38.p14 using MIRAGE2 (Nord et al. 2018). Candidate DCRs were extracted from the MIRAGE alignments using custom scripts and further filtered as described in the result section. From 2037 DCR candidates, we first excluded 237 cases due intron-retention events, and further discarded 229 others for which we noted improper protein mapping (either not starting in a methionine or not ending with a stop codon). Next, 38 DCRs were excluded as at least one isoform involved did not meet our 80% identity threshold, and 17 more had isoform mapping ambiguously assigned to different chromosomes. Sixty-one more cases were further removed as they were redundant, overlapping larger regions. Finally, after a first pass of our classification pipeline, 24 cases were removed following manual inspection (false positives) and 24 others (representing 12 genes) were classified as complex cases and discarded from the main bioinformatic analyses (see Results).

### Canonical vs Alternative frame assignment

Whenever required (differential frame usage in transcriptome and DCR peptide structure prediction) we attempted to assign a canonical or alternative frame to the isoform involved in a given DCR. We used the frame used by the canonical isoform as reported by UniprotKB/Swiss-Prot to report this frame as canonical. In the rare cases where the canonical isoform was not involved in the DCR, the term “canonical” was assigned to the most frequently used among isoforms. In case of tie, no frame assignment was given.

### Susceptibility to nonsense mediated decay (NMD)

The susceptibility of DCR isoforms to NMD was evaluated by examining the distance between the stop codon of each isoform to the last exon-exon junction of the gene model. Gene models were built for each DCR by combining all the exons of the isoforms involved in the gene. Under the assumption that the gene model represents the gene pre-mRNA and NMD is triggered when a premature stop codon (PTC) is located >50bp upstream of the last exon-exon junction, we were able to classify isoforms in three categories: (i) *NMD* (as described by the rule above), (ii) *DCR exons not C-terminal* in any isoform (i.e. DCR is not associated with protein truncation) and (iii) *not enough evidence*, if the putative PTC is located <50bp from the final observed exon-exon junction. This final class is uncertain because we are mapping protein sequences, and may therefore be unaware of 3’UTR exons that could possibly activate the NMD condition (UTRs can span multiple exons).

### DCR conservation

#### Conservation of the DCE status between human and mice

We identified conservation of the DCR status (i.e. whether a gene region contains multiple open reading frames) between human and mice by transferring the coordinates of each human dual coding region (in hg38.p14 coordinates) to the mouse reference genome (mm39) using LiftOver from the UCSC genome browser website (https://genome-euro.ucsc.edu/cgi-bin/hgLiftOver). For DCRs spanning multiple exons, each exon was treated as a separate genomic interval. For each corresponding mouse genomic coordinate, we extracted the DNA sequence from the mm39 assembly using bedtools v2.31.0 (Quinlan 2014) and predicted all sense open reading frames (ORF) with the esl-translate command from easel v.0.48 (Eddy 2011). We counted a DCR region as conserved in mice if at least two different frames could contain an ORF over the full length of the mouse region mapped from the human DCR, allowing for two codons on both 5’ and 3’ ends to be missed by the ORFs. We further produced two control datasets. The first one consists of 100,000 randomly generated intervals within mouse orthologs of non-DCR genes, with length distribution matching that of the human DCRs. These random intervals were used to predict whether at least two ORFs can be produced, with the same full-length requirements as above. Finally, a second control set was produced by randomly shuffling codons within 258,819 mouse orthologs of non-DCR genes (exons ≥21nt and ≤500nt, excluding unplaced scaffolds) and following the same full-length rule (**Figure 2B**). For each dataset, the proportion of regions with 2 or 3 ORFs was reported in bins of 3bp for regions *i* < 100 bp, after which the bin size was dynamically adjusted using 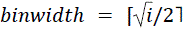.

#### DCR sequence evolution across 100 vertebrates (PhyloP 100-ways)

Conservation of the DCR sequences was addressed by calculating the per-exon mean PhyloP-100 Ways conservation score (which captures per-nucleotide conservation levels across 100 vertebrate genomes) for three category of exons: (i) exons overlapping a dual coding regions (DCR_exon, D), (ii) exons outside of the DCR in a DCR genes (nDCR_exon_DCR_gene, D*) and (iii) exons from non-DCR genes (nDCR_exon_nDCR_gene, ND). The per-base phyloP100-Ways scores for hg38 were downloaded from the UCSC genome browser (https://hgdownload.soe.ucsc.edu/goldenPath/hg38/phyloP100way/hg38.phyloP100way.bw). Significance was assessed by pairwise t-tests within across categories, applying Bonferroni correction to the *P*-values, using the package *ggpubr* version 0.6.0 (Kassambara 2023). To control for potential confounding by length, we matched the two non-DCR categories (D* and ND) to the DCR_exon (D) category by resampling without replacement within the same length distribution as the observed DCR interval. To summarize the results at the gene level, we further calculated the mean phyloP100-Ways score for each DCR (which can overlap multiple exons) and identified outliers representing the most conserved and most diversified DCR genes, using as thresholds the first and 99th centiles of the non-DCR mean phyloP score distribution.

### DCR expression across GTEx tissues

#### Tissue Expression Data

We selected up to 10 random SRA accessions from 53 tissues (only 6 samples from ectocervix and fallopian tubes, and 5 from the endocervix were available) provided by the GTEx consortium (GTEx Consortium 2020) (Table S3). In addition, we collected mouse tissue expression data from (Li et al. 2017), retrieving 72 RNA-seq runs from 18 tissues (between 2 and 8 samples per tissue, as available). We further selected a subset of 14 tissues to match the human GTEx samples (Table S4, Table S5).

#### Frame-specific expression quantification with Tallyman

In order to quantify frame-specific expression of DCR isoforms across human GTEx tissues, we first established a fasta library of 32 bp exon-exon junctions, consisting of 16 bp from the upstream or downstream non-DCR exon, and 16 bp from the DCR 5’ or 3’ end, respectively. This way, each sequence in our library is specific to a particular exon-exon junction in a given frame. Quantification was achieved by reporting the number of RNA-seq reads exactly matching each 32 bp junction. Forward and reverse paired-end reads were pooled in a single file before quantification. Quantification was performed using a custom-built program written in Rust, called Tallyman, available on Github: https://github.com/TravisWheelerLab/tallyman, which produces counts of perfect matches with high speed and without requiring expensive preprocessing. We used the same approach for mouse data, designing a 32bp frame-specific exon-exon junction for 12 DCR with orthologs in mice (Table S6). The sequence of each exon-exon junction was manually created using RefSeq or TOGA genomic annotation available for mm39 in the UCSC genome browser (https://genome.ucsc.edu/cgi-bin/hgTracks?db=mm39).

#### Differential frame x tissue expression analysis

Each analyzed DCR includes up to two junctions: upstream (5’) and/or downstream (3’) of the DCR. To be included in the analysis, a DCR must have at least one end with exon-exon junctions in two or three competing frames. The set of competing exon-exon junctions at a given end of the DCR was defined as a testing unit (e.g. DCR523-upstream). DCR with no distinguishable exon-exon junction between isoforms using different frames were discarded; these include cases where the DCRs are caused by an alternative start site and no discriminatory exon-exon junction is available downstream, or cases where the connecting exons are shorter than 16nt and thus cannot be quantified with our pipeline. We then applied the following procedure for each testing unit. First, if multiple distinct junctions exist at the same end for a given frame (which occurs if two isoforms present distinct junctions connecting to the same DCR frame), the raw counts were summed for the frame. For each testing unit, a matrix with dimensions CxN (C = number of frames; N = number of tissues) was filtered to include only tissues with an average raw count (summed across all junctions in all frames) per sample is at least 2. Furthermore, tissues were retained only if at least one of the frames exhibited a proportion at least 40% greater than the other frame (e.g. a 70/30 split of two frames). Testing proceeded if these conditions were met and at least two tissues remained in the matrix. Such a filter ensured that we only compared tissues with substantial usage differences between DCR frames. Next, goodness-of-fit binomial (for two frames) or multinomial (for three frames) tests were performed between pairs of tissues. In these tests, one tissue served as the “expected” value, while the other represented the “observed” value. The test evaluates whether the observed distribution significantly differs from the expected distribution in a two-sided manner. The smallest *P*-value across all tests, following Bonferroni correction, was retained per testing unit and was deemed significant if *P* < 0.05. For two frames, the R base function ‘binom.test()’ was used. For three frames, if the total number of events (numEvents) was ≤1,000,000, the ‘ExactMultinomialTest()’ function from the EMT package (Menzel 2024) was applied. Otherwise, the ‘MonteCarloMultinomialTest()’ function from the EMT package was used with ntrial = numEvents * 10 and atOnce = 1000000. numEvents was calculated by 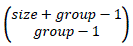, with size set to the total counts in a given tissue, and group set to the total number of frames competing in the junction (n = 3). Finally, a filter was applied to the results in order to identify disordinal tissue-by-frame interactions, i.e. DCR junctions for which the frame with the most count is different between at least two tissues.

#### CPM-TMM normalization

Though our differential frame usage test relies on raw counts, figure representation displays normalized expression in count per millions (CPM) using each library size and trimmed mean of M-values (TMM) scaling factor as implemented in the R package edgeR (Chen et al. 2014).

### Gene Ontology (GO) terms enrichment analysis

Gene Ontology (GO) terms enrichment analysis were performed with the PANTHER classification system (Mi et al. 2019), accessed online at https://geneontology.org/.

### DCR peptide structure prediction

We investigated protein structure in DCRs by selecting pairs of isoforms representative of each frame in a subset of DCRs where the dual coding property does not cause an N- or C- terminal truncation. We thus selected a total of 56 pairs of isoforms by choosing those sharing the most non-DCR exons. Each isoform protein sequence was submitted to the AlphaFold3 (Abramson et al. 2024) webserver. The corresponding prediction, in CIF format, was parsed with DSSP (Hekkelman et al. 2025) to extract the secondary structure classification of each amino-acid. Within the DCR region of each isoform, we retained the DSSP structure classification only if the AlphaFold3 model had a plDDT >= 70.

## Supporting information

Supplementary Figures (1 through 5)

## Data Availability Statement

All presented Datasets, custom programs and analysis scripts are publicly available at https://github.com/TravisWheelerLab/DCR-project

## Acknowledgements

This work was supported by the National Human Genome Research Institute at the National Institute of Health [grant number R21HG012283]. The authors would like to thank Tomi Suomi and Laura Elo for their advice regarding the statistical analysis of expression data.

## Notes

https://github.com/TravisWheelerLab/DCR-project

### Competing Interest Statement

The authors have declared no competing interest.

### Summary of Updates

Update link to Github repository Update First author Initials

## References

1. Abramson J et al. 2024. Accurate structure prediction of biomolecular interactions with AlphaFold 3. Nature. 630:493–500.

2. Ahmed ZM et al. 2008. Mutations of LRTOMT, a fusion gene with alternative reading frames, cause nonsyndromic deafness in humans. Nat. Genet. 40:1335–1340.

3. Alfonso-Gonzalez C, Hilgers V. 2024. (Alternative) transcription start sites as regulators of RNA processing. Trends Cell Biol. 34:1018–1028.

4. Blum M et al. 2025. InterPro: the protein sequence classification resource in 2025. Nucleic Acids Res. 53:D444–D456.

5. Burset M, Seledtsov IA, Solovyev VV. 2000. Analysis of canonical and non-canonical splice sites in mammalian genomes. Nucleic Acids Res. 28:4364–4375.

6. Chen Y, Lun ATL, Smyth GK. 2014. Differential expression analysis of complex RNA-seq experiments using edgeR. In: Statistical Analysis of Next Generation Sequencing Data. Springer International Publishing: Cham pp. 51–74.

7. Chung WY, Wadhawan S, Szklarczyk R, Pond SK, Nekrutenko A. 2007. A first look at ARFome: Dual-coding genes in mammalian genomes. PLoS Comput. Biol. 3:0855–0861.

8. Cozier GE, Carlton J, Bouyoucef D, Cullen PJ. 2004. Membrane targeting by pleckstrin homology domains. Curr. Top. Microbiol. Immunol. 282:49–88.

9. Eddy SR. 2011. Accelerated profile HMM searches. PLoS Comput. Biol. 7:e1002195.

10. de la Fuente R, Díaz-Villanueva W, Arnau V, Moya A. 2025. Alternative splicing across the tree of life. Elife. 13. doi: 10.7554/elife.94802.3.

11. GTEx Consortium. 2020. The GTEx Consortium atlas of genetic regulatory effects across human tissues. Science. 369:1318–1330.

12. Hameed M, Orrell RW, Cobbold M, Goldspink G, Harridge SDR. 2003. Expression of IGF-I splice variants in young and old human skeletal muscle after high resistance exercise. J. Physiol. 547:247–254.

13. Hekkelman ML, Salmoral DÁ, Perrakis A, Joosten RP. 2025. DSSP 4: FAIR annotation of protein secondary structure. Protein Sci. 34:e70208.

14. Hug N, Longman D, Cáceres JF. 2016. Mechanism and regulation of the nonsense-mediated decay pathway. Nucleic Acids Res. 44:1483–1495.

15. Kassambara A. 2023. ggpubr: ‘ggplot2’ Based Publication Ready Plots. https://rpkgs.datanovia.com/ggpubr/.

16. Klemke M, Kehlenbach RH, Huttner WB. 2001. Two overlapping reading frames in a single exon encode interacting proteins - A novel way of gene usage. EMBO J. 20:3849–3860.

17. Kovacs E, Tompa P, Liliom K, Kalmar L. 2010. Dual coding in alternative reading frames correlates with intrinsic protein disorder. Proc. Natl. Acad. Sci. U. S. A. 107:5429–5434.

18. Kozak M. 2001. Extensively overlapping reading frames in a second mammalian gene. EMBO Rep. 2:768–769.

19. Lee S, Aubee JI, Lai EC. 2023. Regulation of alternative splicing and polyadenylation in neurons. Life Sci. Alliance. 6. doi: 10.26508/lsa.202302000.

20. Lewis BP, Green RE, Brenner SE. 2003. Evidence for the widespread coupling of alternative splicing and nonsense-mediated mRNA decay in humans. Proc. Natl. Acad. Sci. U. S. A. 100:189–192.

21. Liang H, Landweber LF. 2006. A genome-wide study of dual coding regions in human alternatively spliced genes. Genome Res. 16:190–196.

22. Li B et al. 2017. A comprehensive mouse transcriptomic BodyMap across 17 tissues by RNA-seq. Sci. Rep. 7:4200.

23. Linares AJ et al. 2015. The splicing regulator PTBP1 controls the activity of the transcription factor Pbx1 during neuronal differentiation. Elife. 4:e09268.

24. Ling JP et al. 2016. PTBP1 and PTBP2 repress nonconserved cryptic exons. Cell Rep. 17:104–113.

25. Lin MF et al. 2011. Locating protein-coding sequences under selection for additional, overlapping functions in 29 mammalian genomes. Genome Res. 21:1916–1928.

26. Makeyev EV, Zhang J, Carrasco MA, Maniatis T. 2007. The MicroRNA miR-124 promotes neuronal differentiation by triggering brain-specific alternative pre-mRNA splicing. Mol. Cell. 27:435–448.

27. Marasco LE, Kornblihtt AR. 2023. The physiology of alternative splicing. Nat. Rev. Mol. Cell Biol. 24:242–254.

28. Mauger O, Scheiffele P. 2017. Beyond proteome diversity: alternative splicing as a regulator of neuronal transcript dynamics. Curr. Opin. Neurobiol. 45:162–168.

29. Mayr C. 2017. Regulation by 3’-untranslated regions. Annu. Rev. Genet. 51:171–194.

30. Mayr C. 2019. What are 3’ UTRs doing? Cold Spring Harb. Perspect. Biol. 11:a034728.

31. Menzel U. 2024. Exact Multinomial Test: Goodness-of-Fit Test for Discrete Multivariate Data. https://cran.r-project.org/web/packages/EMT.

32. Michel AM et al. 2012. Observation of dually decoded regions of the human genome using ribosome profiling data. Genome Res. 22:2219–2229.

33. Mi H et al. 2019. Protocol Update for large-scale genome and gene function analysis with the PANTHER classification system (v.14.0). Nat. Protoc. 14:703–721.

34. Nagy E, Maquat LE. 1998. A rule for termination-codon position within intron-containing genes: when nonsense affects RNA abundance. Trends Biochem. Sci. 23:198–199.

35. Najar CFBA et al. 2025. Genetic and functional analysis of unproductive splicing using LeafCutter2. Genomics. https://www.biorxiv.org/content/10.1101/2025.04.06.646893v1.full.

36. Nord A, Hornbeck P, Carey K, Wheeler T. 2018. Splice-Aware Multiple Sequence Alignment of Protein Isoforms. ACM-BCB 2018 - Proceedings of the 2018 ACM International Conference on Bioinformatics, Computational Biology, and Health Informatics. 18:200–210.

37. Perez G et al. 2025. The UCSC Genome Browser database: 2025 update. Nucleic Acids Res. 53:D1243–D1249.

38. Pollard KS, Hubisz MJ, Rosenbloom KR, Siepel A. 2010. Detection of nonneutral substitution rates on mammalian phylogenies. Genome Res. 20:110–121.

39. Poulin F, Brueschke A, Sonenberg N. 2003. Gene Fusion and Overlapping Reading Frames in the Mammalian Genes for 4E-BP3 and MASK. J. Biol. Chem. 278:52290–52297.

40. Preussner M et al. 2020. Splicing-accessible coding 3’UTRs control protein stability and interaction networks. Genome Biol. 21:186.

41. Quinlan AR. 2014. BEDTools: The Swiss-Army Tool for Genome Feature Analysis. Curr. Protoc. Bioinformatics. 47:11.12.1–34.

42. Ravichandran Y et al. 2024. The distinct localization of CDC42 isoforms is responsible for their specific functions during migration. J. Cell Biol. 223. doi: 10.1083/jcb.202004092.

43. R Core Team. 2023. R: A Language and Environment for Statistical Computing. https://www.R-project.org/.

44. Ribrioux S, Brüngger A, Baumgarten B, Seuwen K, John MR. 2008. Bioinformatics prediction of overlapping frameshifted translation products in mammalian transcripts. BMC Genomics. 9:1–16.

45. Rodriguez JM, Pozo F, di Domenico T, Vazquez J, Tress ML. 2020. An analysis of tissue-specific alternative splicing at the protein level. PLoS Comput. Biol. 16:e1008287.

46. Romero PR et al. 2006. Alternative splicing in concert with protein intrinsic disorder enables increased functional diversity in multicellular organisms. Proc. Natl. Acad. Sci. U. S. A. 103:8390–8395.

47. Schwarz DA et al. 2000. Characterization of gamma-aminobutyric acid receptor GABAB(1e), a GABAB(1) splice variant encoding a truncated receptor. J. Biol. Chem. 275:32174–32181.

48. Scorilas A et al. 2001. Molecular cloning, physical mapping, and expression analysis of a novel gene, BCL2L12, encoding a proline-rich protein with a highly conserved BH2 domain of the Bcl-2 family. Genomics. 72:217–221.

49. Sharpless NE, DePinho RA. 1999. The INK4A/ARF locus and its two gene products. Curr. Opin. Genet. Dev. 9:22–30.

50. Szklarczyk R, Heringa J, Pond SK, Nekrutenko A. 2007. Rapid asymmetric evolution of a dual-coding tumor suppressor INK4a/ARF locus contradicts its function. Proc. Natl. Acad. Sci. U. S. A. 104:12807–12812.

51. UniProt Consortium. 2025. UniProt: The universal protein knowledgebase in 2025. Nucleic Acids Res. 53:D609–D617.

52. Vanderperre B et al. 2013. Direct Detection of Alternative Open Reading Frames Translation Products in Human Significantly Expands the Proteome. PLoS One. 8:e70698–e70698.

53. Vuong JK, Ergin V, Chen L, Zheng S. 2022. Multilayered regulations of alternative splicing, NMD, and protein stability control temporal induction and tissue-specific expression of TRIM46 during axon formation. Nat. Commun. 13:2081.

54. Weise A et al. 2010. Alternative splicing of Tcf7l2 transcripts generates protein variants with differential promoter-binding and transcriptional activation properties at Wnt/beta-catenin targets. Nucleic Acids Res. 38:1964–1981.

55. Xu H et al. 2010. Length of the ORF, position of the first AUG and the Kozak motif are important factors in potential dual-coding transcripts. Cell Res. 20:445–457.

56. Yoshida H, Matsui T, Yamamoto A, Okada T, Mori K. 2001. XBP1 mRNA is induced by ATF6 and spliced by IRE1 in response to ER stress to produce a highly active transcription factor. Cell. 107:881–891.

57. Zhao FQ, Zheng Y, Dong B, Oka T. 2004. Cloning, genomic organization, expression, and effect on β-casein promoter activity of a novel isoform of the mouse Oct-1 transcription factor. Gene. 326:175–187.

