## Supplementary Figures (1 through 5) for "Alternatively Spliced Dual-Coding Regions Contribute to the Human Gene Regulatory Program"

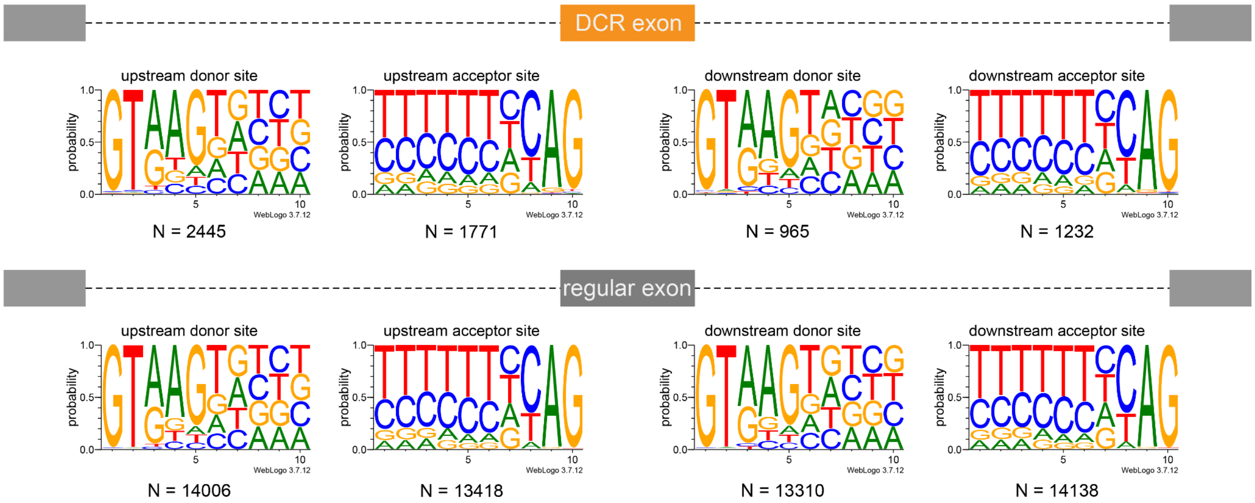


**Figure S1.** Logo representation of the exon-exon junctions motifs upstream and downstream DCRs-overlapping and control ("regular") exons.

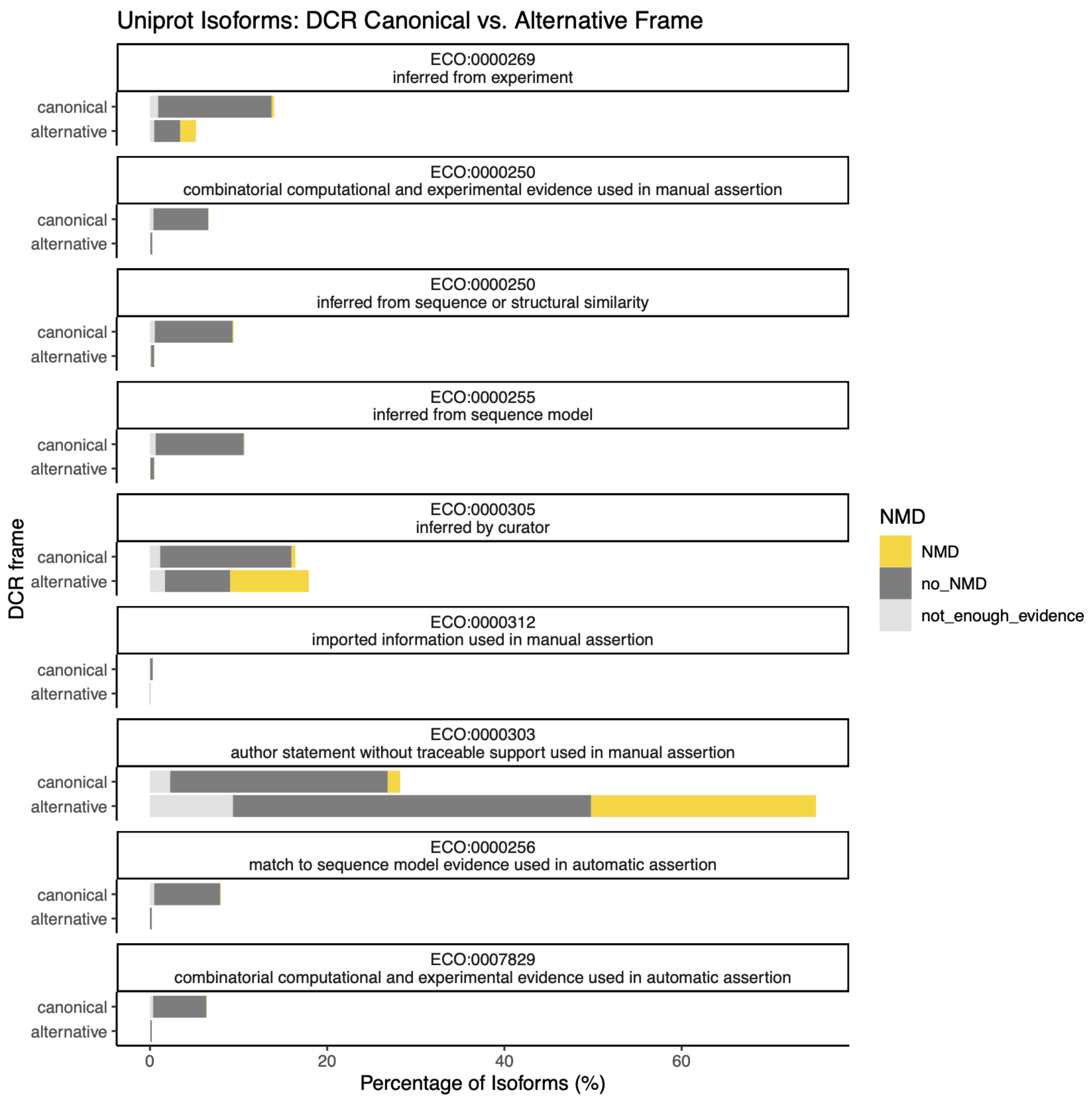


**Figure S2.** Distribution of the UniprotKB/Swiss-Prot annotation codes according to the isoform canonical or alternative status determined by Uniprot and NMD susceptibility classification determined by our analysis.

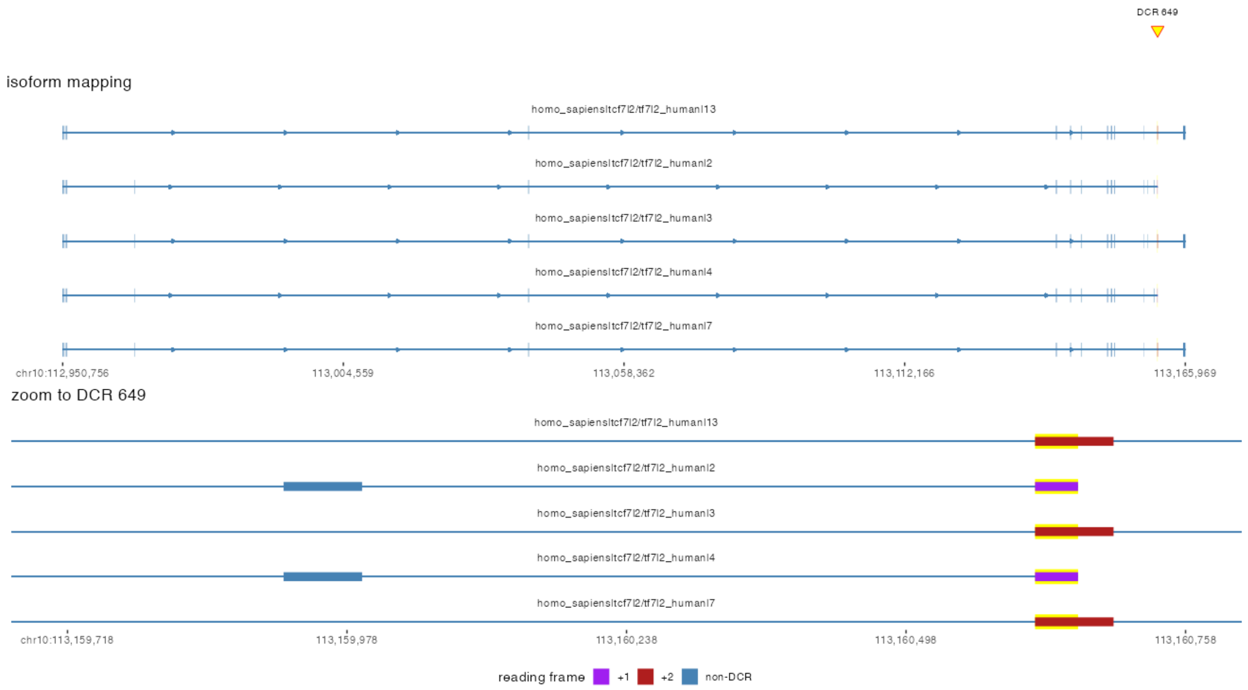


**Figure S3.** Gene model and zoom to the DCR region 649 for the gene *TCF7L2*. Note: no canonical frame can be assigned as the Uniprot canonical isoform (#1) is not involved in the DCR. Isoforms 3, 7 and 13 are representatives of isoforms E according to Weisse et al. 2010.

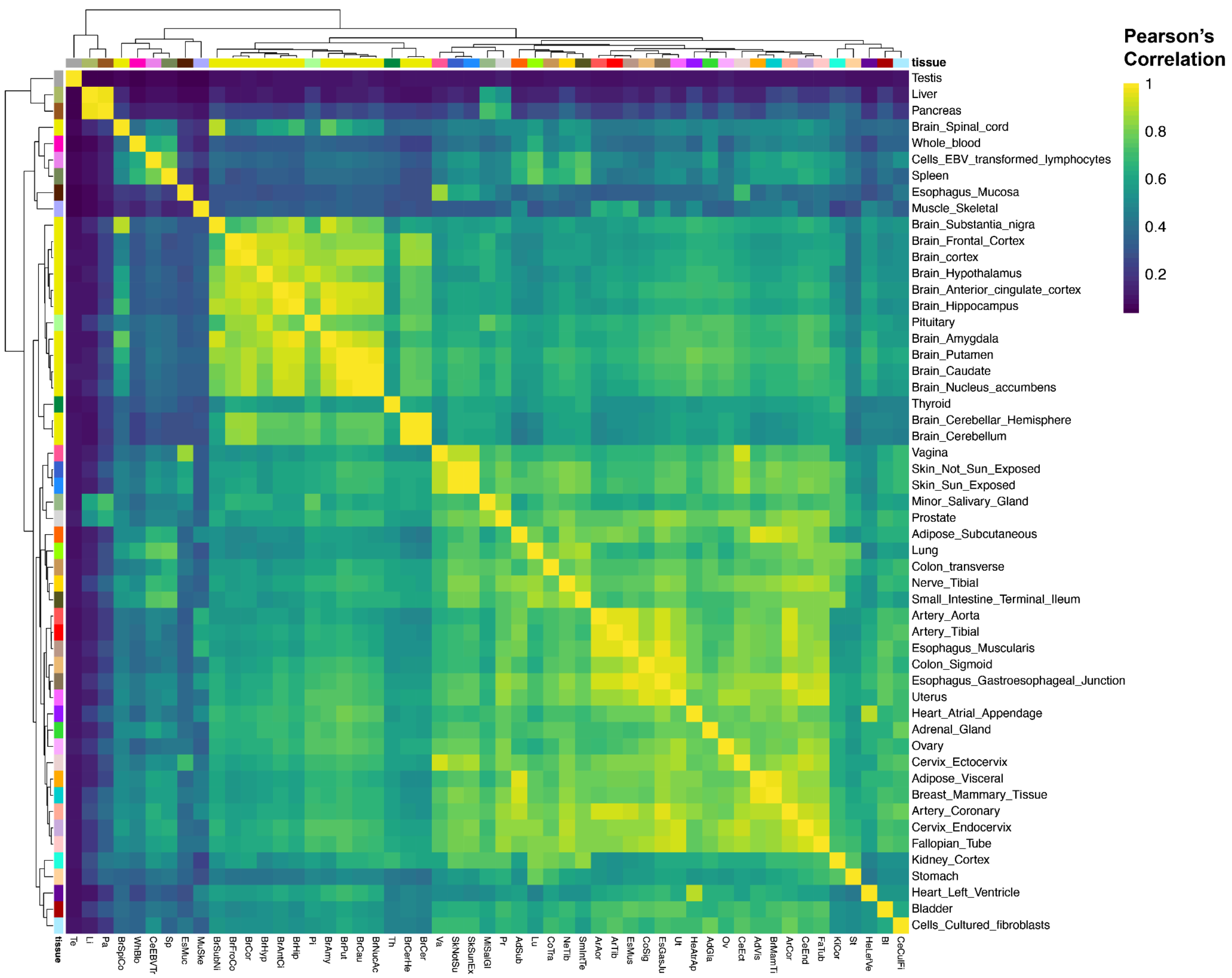


**Figure S4.** Pearson's correlation between 53 GTEx tissues based on the quantification of exon-exon junction (EEJs) at DCRs, using EEJs with an average of 2 reads per sample in at least 2 tissues. Row and Columns: GTEx tissues. Dendograms: hierarchical clustering (method = "complete", distance = "Euclidian").

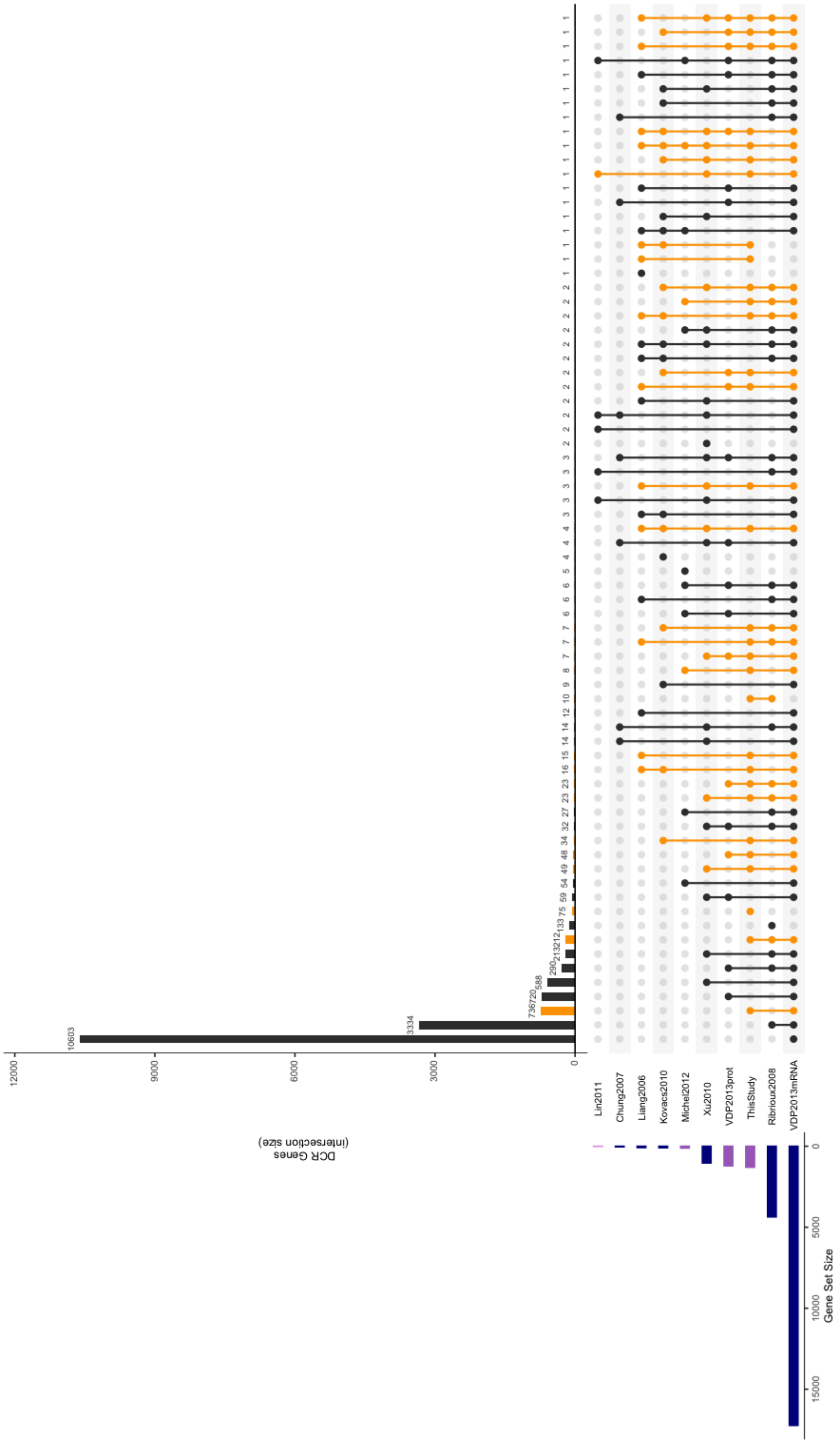


**Figure S5.** Upset plot comparison of DCR genes reported across genome-wide studies. Intersections with this study are highlighted in yellow. Studies based on protein evidence are highlighted in purple.
